# Introductions and early spread of SARS-CoV-2 in France

**DOI:** 10.1101/2020.04.24.059576

**Authors:** Fabiana Gámbaro, Sylvie Behillil, Artem Baidaliuk, Flora Donati, Mélanie Albert, Andreea Alexandru, Maud Vanpeene, Méline Bizard, Angela Brisebarre, Marion Barbet, Fawzi Derrar, Sylvie van der Werf, Vincent Enouf, Etienne Simon-Loriere

## Abstract

Following the emergence of coronavirus disease (COVID-19) in Wuhan, China in December 2019, specific COVID-19 surveillance was launched in France on January 10, 2020. Two weeks later, the first three imported cases of COVID-19 into Europe were diagnosed in France. We sequenced 97 severe acute respiratory syndrome coronavirus 2 (SARS-CoV-2) genomes from samples collected between January 24 and March 24, 2020 from infected patients in France. Phylogenetic analysis identified several early independent SARS-CoV-2 introductions without local transmission, highlighting the efficacy of the measures taken to prevent virus spread from symptomatic cases. In parallel, our genomic data reveals the later predominant circulation of a major clade in many French regions, and implies local circulation of the virus in undocumented infections prior to the wave of COVID-19 cases. This study emphasizes the importance of continuous and geographically broad genomic sequencing and calls for further efforts with inclusion of asymptomatic infections.

## Introduction

Severe acute respiratory syndrome coronavirus 2 (SARS-CoV-2) was identified as the cause of an outbreak of severe respiratory infections in Wuhan, China in December 2019 (Zhu, Zhang et al. 2020). Although Chinese authorities implemented strict quarantine measures in Wuhan and surrounding areas, this emerging virus has rapidly spread across the globe, and the World Health Organization (WHO) declared a pandemic of coronavirus disease 2019 (COVID-19) on March 11, 2020.

Strengthened surveillance of COVID-19 cases was implemented in France on January 10, 2020, with the objective of identifying imported cases early to prevent secondary transmission in the community, and the National Reference Center for Respiratory Viruses (NRC) hosted at Institut Pasteur identified the first cases in Europe. With the extension of the epidemic, identification of SARS-CoV-2 cases was shared with the NRC associated laboratory in Lyon and then extended to additional first line hospital laboratories, with the NRC at Institut Pasteur focusing on the Northern part of France, including the densely populated capital. Screening and sampling for SARS-CoV-2 was targeted towards patients who had symptoms (fever and/or respiratory problems) or had travel history to risk zones of infection. With the spread of the virus, it became clear that clinical characteristics of COVID-19 patients vary greatly (Guan, Ni et al. 2020, Onder, Rezza et al. 2020) with a proportion of asymptomatic infections or mild disease cases (Li, Pei et al. 2020).

Viral genomics coupled with modern surveillance systems is transforming the way we respond to emerging infectious diseases (Gardy and Loman 2018, Ladner, Grubaugh et al. 2019). Realtime genomic epidemiology data has proven to be useful to reconstruct outbreak dynamics: from virus identification to understanding the factors contributing towards global spread (Grubaugh, Ladner et al. 2019). Here, we sequenced SARS-CoV-2 genomes from clinical cases sampled since the beginning of the syndromic surveillance in France. We used the newly generated genomes to investigate the origins of SARS-CoV-2 lineages circulating in Northern France and better understand its spread.

## Results/Discussion

We generated complete SARS-CoV-2 genome sequences from nasopharyngeal or sputum samples addressed to the National Reference Center for Respiratory Viruses at the Institut Pasteur in Paris as part of the ongoing surveillance (Figure 1A). We combined the 97 SARS-CoV-2 genomes from France and 3 from Algeria generated here with 338 sequences published or freely available from the GISAID database, and performed a phylogenetic analysis focusing on the initial introductions and spread of the virus in France.

**Figure 1.**
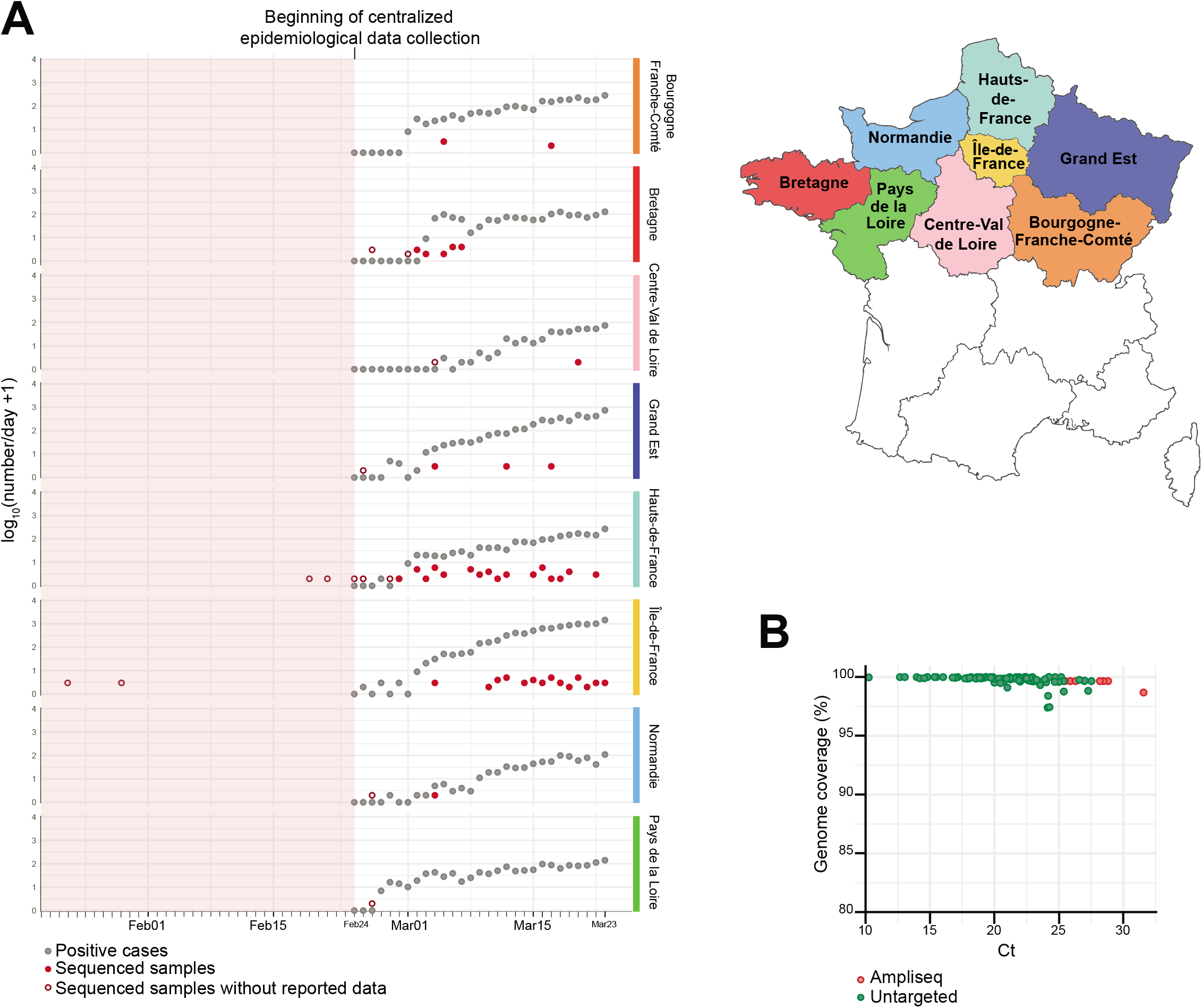
SARS-CoV-2 genome sequencing effort in the Northern French regions in a rapidly growing pandemic. **A.** The plot represents the numbers of daily sequenced genomes in this study (red filled or hollow circles) overlaid with the number of reported positive cases (grey circles) obtained from Santé Publique France (www.santepubliquefrance.fr). Hollow circles indicate samples obtained on dates with zero reported positive cases. The data are shown separately for each region of Northern France as indicated on the map on the right. **B.** Percentage of SARS-CoV-2 genome coverage obtained in relation to the Ct values for the 97 genomes reported here. Colors indicate sequencing approach: untargeted metagenomics (green) or amplicon-based sequencing (red).

### Early introductions do not appear to have resulted in local transmission

Our analysis indicates that the quarantine imposed on the initial COVID-19 cases in France appears to have prevented local transmission. The first European cases sampled on January 24, 2020 (IDF0372 and IDF0373 from Île-de-France, described in (Lescure, Bouadma et al. 2020) were direct imports from Hubei, China, and the genomes fall accordingly near the base of the tree, within clade V, according to GISAID nomenclature (Figure 2, Figure 3A). These identical genomes both harbor a V367F (G22661T) mutation in the receptor binding domain of the Spike, not observed in other genomes. Similarly, IDF0515 corresponds to a traveler from Hubei, China. This basal genome falls outside of the three major GISAID proposed clades V, G, and S (Figure 2, Figure 3A), but carries the G11083T mutation associated with putative lineage V1 (Figure 3A, Figure S2), suggesting convergent evolution or more likely a reversion of the V-clade defining G26144T change. Subsequent early cases in the West or East of France (B2334/B2340, clade V and GE1583, clade S), all with recent history of travel to Italy, add to the genomic diversity of viruses from Northern Italy, but also do not appear to have seeded local transmission with the current sampling (Figure 2).

**Figure 2.**
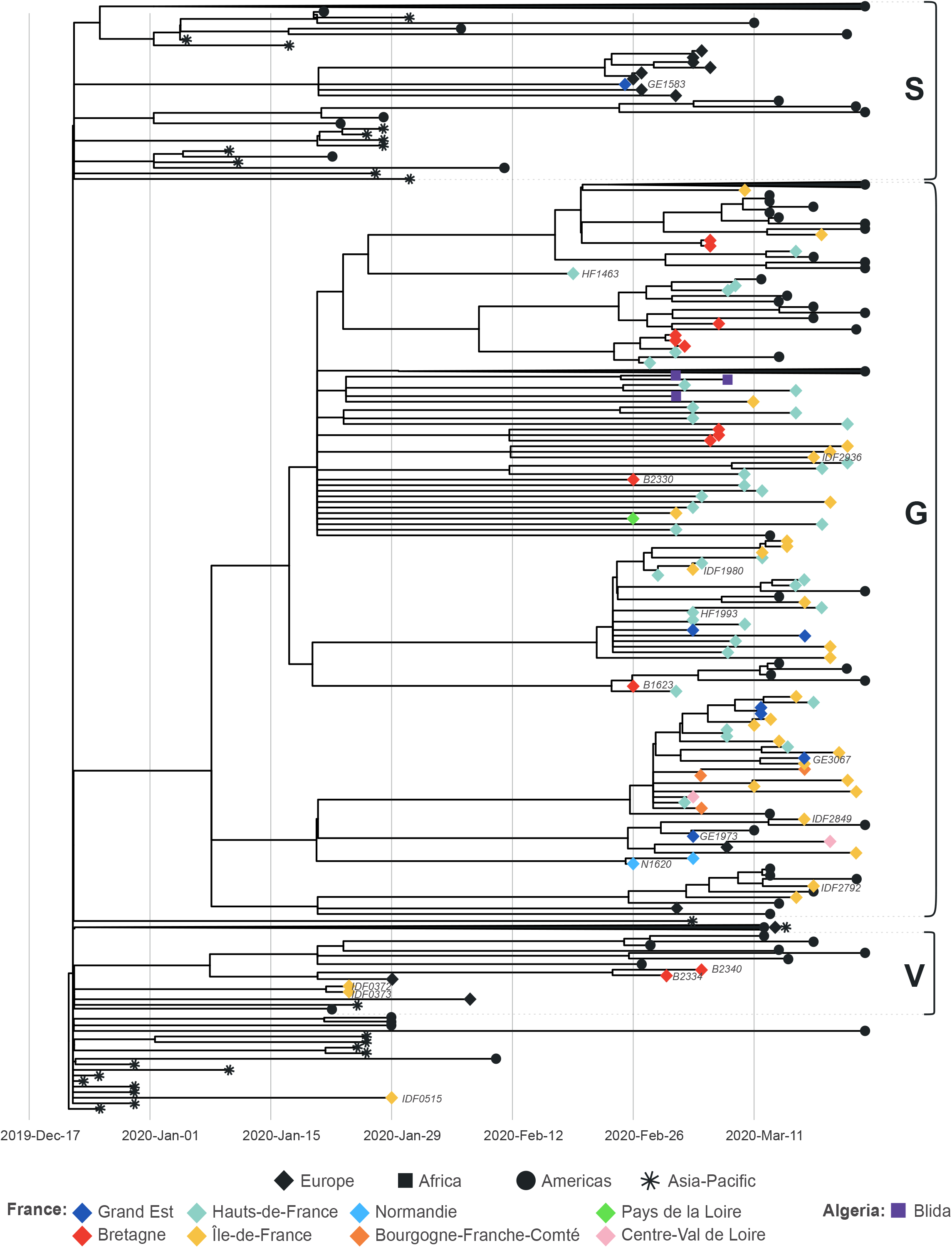
Northern France sequences from early introductions and currently circulating lineages. Time calibrated tree of 438 SARS-CoV-2 sequences including Northern France, Algeria and publicly available global sequences. The tree is rooted using the reference strain Wuhan/Hu-1/2019n (MN908947). The tips of the tree are shaped and colored according to sampling location. Branch lengths are proportional to the time span from the sampling date to the inferred date of the most recent common ancestor. The three major clades according to GISAID nomenclature are indicated. Strain names of the sequences discussed in this study are indicated next to the corresponding tips.

### The current outbreak lineages

All other sequences from Northern France fall in clade G (defined by a single non-synonymous mutation, D614G (A23403G) in the Spike, Figure 2), and this includes sequences captured during the steep increase of reported cases in many strongly affected regions (Figure 1). While a more thorough sampling will be needed to confirm this observation, it suggests that, unlike what is observed for many other European countries (Gudbjartsson, Helgason et al. 2020, Zehender, Lai et al. 2020), the French outbreak has been mainly seeded by one or several variants of this clade. This clade can be further classified into lineages, albeit supported again by only 1 to 3 substitutions (putatively named G1, G2, G3, G3a, G3b), and the diversity of the sequence from Northern France is spread out, with most regions represented in the different lineages. Several genomes correspond to patients with recent history of travel in Europe (GE3067, N1620, IDF2792), United Arab Emirates (IDF2936), Madagascar (HF1993) or Egypt (B1623, B2330), and might represent additional introductions of the same clade. On the other hand, in lineage G3b, three sequences sampled in Algeria are closely related to sequences from France and likely represent exported cases in light of recent history of travel to France.

The syndromic surveillance allowed to capture one of the earliest representatives of clade G (HF1463, sampled on February 19th) (Figure 2). Importantly, this sequence carries 2 additional mutations compared to the reconstructed ancestral sequence of this clade (Figure 3B). Other sequences sampled weeks later (IDF2849, GE1973) are more basal to the clade, highlighting the complexity and risk of inferences based on 1 or 2 nucleotide substitutions. Because of this, and the scarcity of early sequences in many countries in Europe, country and within-country level phylogeographic estimations for the origin of clade G are also unreliable with the current dataset.

**Figure 3.**
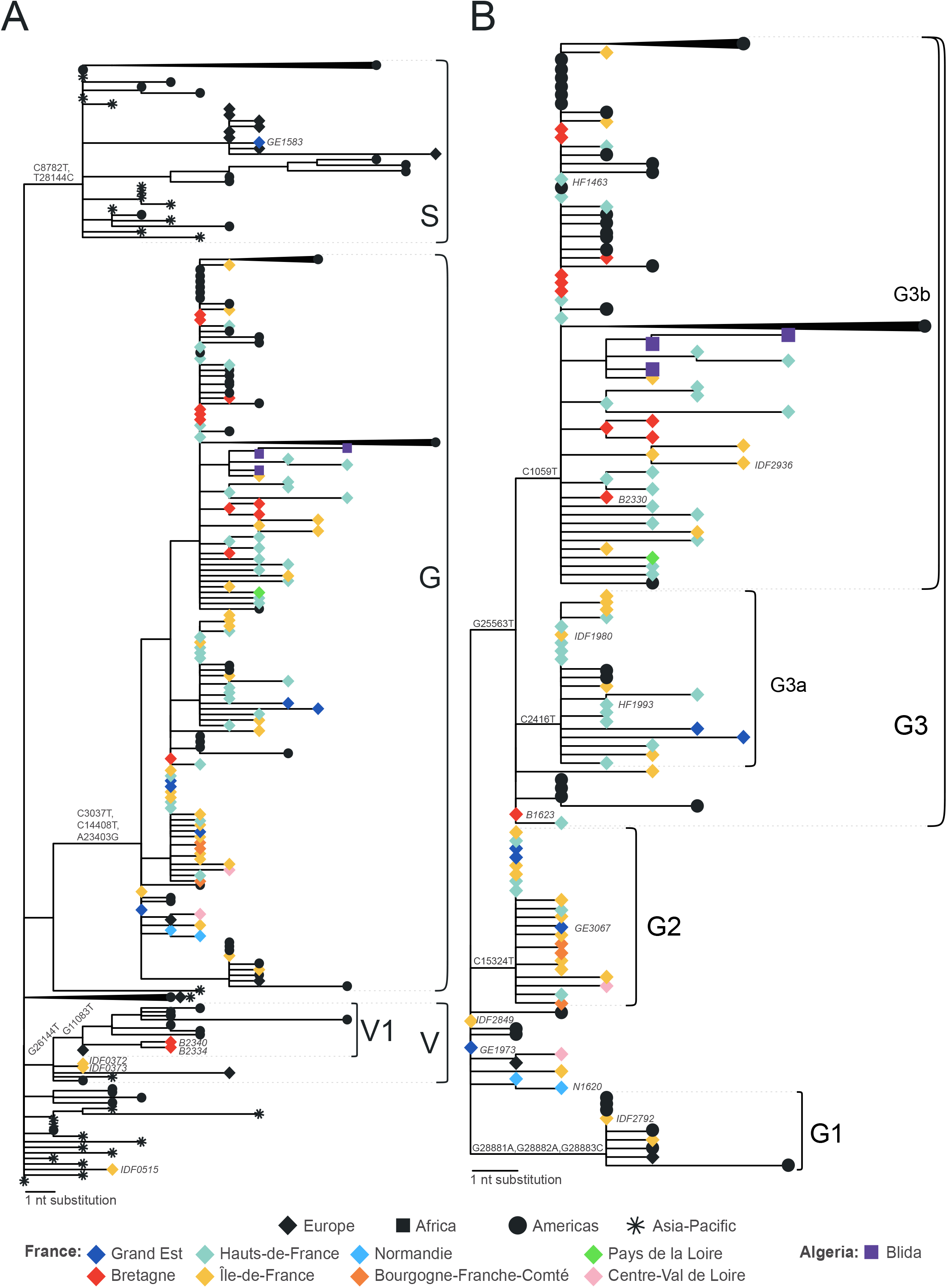
Divergence of SARS-CoV-2 sequences from Northern France. Divergence tree of SARS-CoV-2 genomes including sequences from Northern France with the three major clades (**A**) or only clade G (**B**). The tips of the tree are shaped and colored according to sampling location. Branch lengths are proportional to the number of nucleotide substitutions from the reference and tree root Wuhan/Hu-1/2019 (MN908947). GISAID clades and putative lineages are indicated on the right of each panel. Strain names of the sequences discussed in this study are indicated next to the corresponding tips in italic. Nucleotide substitutions shared among all the sequences of each clade or lineage are indicated next to the corresponding nodes. Some monophyletic lineages are collapsed for ease of representation. A complete tree is shown in Figure S1.

Crucially, while all early symptomatic suspected COVID-19 cases were addressed to the NRC for testing, this was no longer the case as the epidemic developed (Figure 1A). In addition, pauci or asymptomatic cases are scarcely represented here. As the earliest representative of clade G (HF1463) had no history of travel or contact with returning travelers, we can infer that the virus was silently circulating in France in February, a scenario compatible with the large proportion of mild or asymptomatic diseases (Li, Pei et al. 2020), and observations in other European countries (Gudbjartsson, Helgason et al. 2020, Onder, Rezza et al. 2020). While this is also compatible with the time to the most recent common ancestor estimate for clade G (Figure 2), the current sampling clearly prevents reliable inference for the timing of introduction in France.

This study reveals areas for potential improvement of SARS-CoV-2 genomic surveillance in France. Several regions are poorly represented yet, likely due to the heavy burden on hospitals, which were quickly able to perform local testing as the molecular detection tools were rapidly shared by the NRC. Because of this, and of the syndromic-only based surveillance, we likely underestimate the genetic diversity of SARS-CoV-2 circulating in France.

In conclusion, our study sheds light on the origin and diversity of the COVID-19 outbreak in France with insights for Europe, and highlights the challenges of containment measures when a significant proportion of cases are asymptomatic.

## Materials and Methods

### Ethical statement

Samples used in this study were collected as part of approved ongoing surveillance conducted by the National Reference Center for Respiratory Viruses (NRC) at Institut Pasteur (WHO reference laboratory providing confirmatory testing for COVID-19). The investigations were carried out in accordance with the General Data Protection Regulation (Regulation (EU) 2016/679 and Directive 95/46/EC) and the French data protection law (Law 78–17 on 06/01/1978 and Décret 2019–536 on 29/05/2019).

### Sample collection

After reports of severe pneumonia in late December 2019, enhanced surveillance was implemented in France to detect suspected infections. For each suspected case, respiratory samples from the upper respiratory tract (nasopharyngeal swabs or aspirates) and when possible from the lower respiratory tract, were sent to the NRC, to perform SARS-CoV-2-specific real-time RT-PCR. Demographic information, date of illness onset, and travel history, were obtained when possible. A subset of samples were selected according to the viral load and their sampling location in order to have a broad representation across different regions of France.

### Molecular test

RNA extraction was performed with the Extraction NucleoSpin Dx Virus kit (Macherey Nagel). RNA was extracted from 100 μl of specimen, eluted in 100 μl of water and used as a template for RT-qPCR. Samples were tested with a one-step RT-qPCR using three sets of primers as described on the WHO website (https://www.who.int/docs/default-source/coronaviruse/real-time-rt-pcr-assays-for-the-detection-of-sars-cov-2-institut-pasteur-paris.pdf?sfvrsn=3662fcb62).

### Virus Sequencing

Viral genome sequences were generated by two different approaches. The first consisted in direct metagenomic sequencing, which resulted in complete or near complete genome sequences for samples with viral load higher than 1.45×10^4^ viral genome copies/μl, which corresponds to a Ct value of 25.6 with the IP4 primer set (Figure 1B, Supplementary material, Table S3). Briefly, extracted RNA was first treated with Turbo DNase (Ambion) followed by purification using SPRI beads Agencourt RNA clean XP (Beckman Coulter). RNA was converted to double stranded cDNA. Libraries were then prepared using the Nextera XT DNA Library Prep Kit (Illumina) and sequenced on an Illumina NextSeq500 (2×150 cycles) on the Mutualized Platform for Microbiology (P2M) at Institut Pasteur.

For samples with lower viral load, we implemented a highly multiplexed PCR amplicon approach (Quick, Grubaugh et al. 2017) using the ARTIC Network multiplex PCR primers set v1 (https://artic.network/ncov-2019), with modification as suggested in (Kentaro, Tsuyoshi et al. 2020). Synthesized cDNA was used as template and amplicons were generated using two pooled primer mixtures for 35 rounds of amplification. We prepared sequencing libraries using the NEBNext Ultra II DNA Library Prep Kit for Illumina (New England Biolabs) and barcoded with NEBNext Multiplex Oligos for Illumina (Dual Index Primers Set 1) (New England Biolabs). We sequenced prepared libraries on an Illumina MiSeq using MiSeq Reagent Kit v3 (2×300 cycles) at the biomics platform of Institut Pasteur.

### Genome assembly

Raw reads were trimmed using Trimmomatic v0.36 (Bolger, Lohse et al. 2014) to remove Illumina adaptors and low quality reads, as well as primer sequences for samples sequenced with the amplicon-based approach. We assembled reads from all sequencing methods into genomes using Megahit, and also performed direct mapping against reference genome Wuhan/Hu-1/2019 (NCBI Nucleotide – NC_045512, GenBank – MN908947) using the CLC Genomics Suite v5.1.0 (QIAGEN). We then used SAMtools v1.3 to sort the aligned bam files and generate alignment statistics (WysokerA, RuanJ et al.). Aligned reads were manually inspected using Geneious prime v2020.1.2 (2020) (https://www.geneious.com/), and consensus sequences were generated using a minimum of 3X read-depth coverage to make a base call. No genomic deletions were detected in the genomes analyzed.

### Phylogenetic analysis

A set of 100 SARS-CoV-2 sequences generated in this study (97 from France, 3 from Algeria) was complemented with 338 genomes published or freely available sequences on GenBank or the GISAID database. From the latter, only published sequences were chosen (Deng, Gu et al. 2020, Fauver, Petrone et al. 2020) (Supplementary material, Table S2). A total of 438 full genome sequences were analyzed with augur and auspice as implemented in the Nextstrain pipeline (Hadfield, Megill et al. 2018) version from March 20, 2020 (https://github.com/nextstrain/ncov). Within the pipeline, sequences were aligned to the reference Wuhan/Hu-1/2020 strain of SARS-CoV-2 (GenBank accession MN908947) using MAFFT v7.455 (Katoh and Toh 2010). The alignment was visually inspected and sequences from France were subset to analyze shared SNPs. No evidence of recombination was detected using RDP v4.97 (Martin, Murrell et al. 2015). A maximum likelihood phylogenetic tree was built using IQ-TREE v1.5.5 with the GTR model (Kalyaanamoorthy, Minh et al. 2017), after masking 130 and 50 nucleotides from the 5’ and 3’ ends of the alignment, respectively, as well as single nucleotides at positions 18529, 29849, 29851, 29853 as normally set in Nextstrain implementation for SARS-CoV-2. We checked for temporal signal using Tempest v1.5.3 (Rambaut, Lam et al. 2016). The temporal phylogenetic analyses were performed with augur and TreeTime (Sagulenko, Puller et al. 2018), assuming clock rate of 0.0008±0.0004 (SD) substitutions/site/year (Rambaut 2020), coalescent skyline population growth model and the root set on the branch leading to the Wuhan/Hu-1/2020 sequence. The time and divergence trees were visualized with FigTree v1.4.4 (http://tree.bio.ed.ac.uk/software/figtree/). Nucleotide substitutions from the reference sequence that define internal nodes of the tree were extracted from the final Nextstrain build file and annotated on the tree using a custom R script (https://www.R-project.org/) with packages *tidyverse* v1.3.0 (Wickham, Averick et al. 2019) and *jsonlite* v1.6.1 (Ooms 2014). Adobe Illustrator 2020 was used to prepare final tree figures. Sequence metadata (Supplementary material, Table S2), Nextstrain build, and R script is available at https://github.com/Simon-LoriereLab/SARS-CoV-2-France. The phylogeny can be visualized interactively at https://nextstrain.org/community/Simon-LoriereLab/SARS-CoV-2-France. In this study, we used the proposed nomenclature from GISAID to annotate three major clades V, G, and S according to specific single-nucleotide polymorphisms that are shared by all sequences in the clade. Clade defining variants according to GISAID nomenclature are included in Supplementary material, Table S1.

## Data sharing

The assembled SARS-CoV-2 genomes generated in this study were deposited on the GISAID database (https://www.gisaid.org/) as soon as they were generated, accession numbers can be found in Supplementary material, Table S2.

## Acknowledgements

We would like to thank all of the health care workers, public health employees, and scientists involved in the COVID-19 response. We acknowledge the hospital laboratories from the RENAL network in the north of France (list of names in Supplementary material, Table S4). We acknowledge the authors, originating and submitting laboratories of the sequences from GISAID and GenBank (Supplementary material, Table S2). We avoided any direct analysis of genomic data not submitted as part of this paper and used this genomic data only as background. This study has received funding from Institut Pasteur, CNRS, Université de Paris, Santé publique France, the French Government’s Investissement d’Avenir program, Laboratoire d’Excellence “Integrative Biology of Emerging Infectious Diseases” (grant n°ANR-10-LABX-62-IBEID), REACTing (Research & Action Emerging Infectious Diseases), France Génomique (ANR-10-INBS-09-09), IBISA, and the EU grant Recover. ESL acknowledges funding from the INCEPTION program (Investissements d’Avenir grant ANR-16-CONV-0005). We thank Laurence Ma (Biomics Platform, C2RT, Institut Pasteur, Paris, France) for the MiSeq sequencing. This work used the computational and storage services (TARS cluster) provided by the IT department at Institut Pasteur, Paris. FG is part of the Pasteur-Paris University (PPU) International PhD program, BioSPC doctoral school.

## Supplementary figures

**Figure S1.**
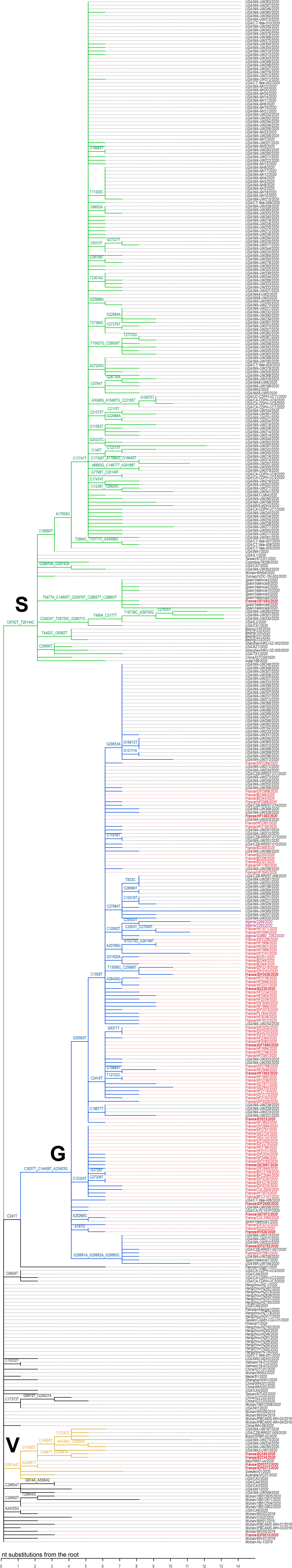
Phylogenetic divergence tree of all labeled SARS-CoV-2 sequences used in this study. Maximum-likelihood tree including all sequences from Northern France, Algerian sequences and publicly available global SARS-CoV-2 sequences, corresponding to the collapsed tree shown in Figure 3 (same ordering as in Figure 2 and Figure 3). GISAID clades are indicated next to the corresponding nodes and branches are colored distinctly. Tips indicate strain names, colored in red for sequences from France and purple for Algeria, and are noted in bold if discussed in this study.

**Figure S2.**
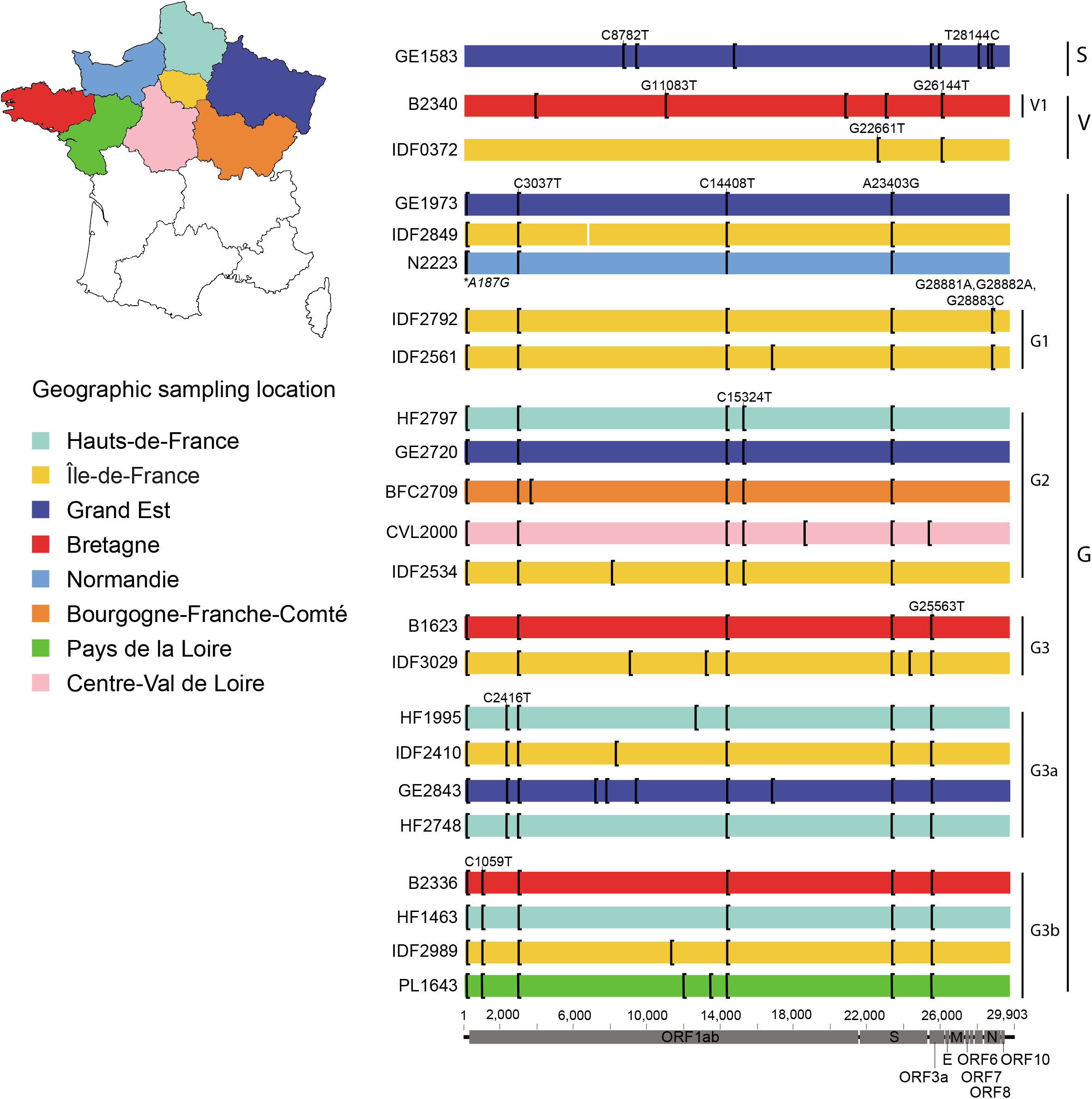
Single-nucleotide polymorphisms representing the diversity among sequences across the regions of Northern France. Multiple sequence alignment of all SARS-CoV-2 genomes sampled across the northern part of France from different clades and lineages. Single nucleotide variants with respect to the reference (MN908947) are shown as black vertical bars and shared substitutions among the sequences of each clade or lineage are annotated. A substitution only found in sequences from Normandie is noted in italic.

## Supplementary material

A single Excel document with multiple sheets, representing tables below.

**Table S1. Clade defining SNPs according to GISAID nomenclature.**

**Table S2. Metadata associated with the sequences used in this study.**

**Table S3. Viral RNA load and genome recovery data.**

**Table S4. List of collaborators in the RENAL network in the north of France.**

